# A tool for the *in vivo* gating of gene expression in neurons using the co-occurrence of neural activity and light

**DOI:** 10.1101/2021.12.05.471336

**Authors:** Adam T. Vogel, Shelley J. Russek

## Abstract

Advancements in genetically based technologies have begun to allow us to better understand the relationships between underlying neural activity and the patterns of measurable behavior that can be reproducibly studied in the laboratory. As this field develops, there are key limitations to the currently available technologies hindering their full potential to deliver meaningful datasets. The limitations which are most critical to advancement of these technologies in behavioral neuroscience are: the temporal resolution at which physiological events can be windowed, the divergent molecular pathways in signal transduction that introduce ambiguity into the output of activity sensors, and the impractical size of the genetic material that requires 3-4 separate AAV vectors to deliver a fully functional system into a cell. To address these limitations and help bring the potential of these types of technologies into better realization, we have engineered a nucleus localized light-sensitive Ca^2+^-dependent gene expression system based on AsLOV_2_ and the downstream responsive element antagonist modulator (DREAM). The design and engineering of each component was performed in such a way to: 1) preserve behaviorally relevant temporal dynamics, 2) preserve signal fidelity appropriate for studying experience-driven neural activity patterns and their relationship to specific animal responses, and 3) have full delivery of the genetic material via a single AAV vector. The system was tested in vitro and subsequently *in vivo* with neural activity induced by Channelrhodopsin and could be used in the future with behaviorally-driven neural activity. To our knowledge this is the first optogenetic tool for the practical use of linking activity-dependent gene activation in response to direct nuclear calcium transduction.

## Main

Neuroscientists have been developing a wide range of new tools to study the patterns of neural activity underlying behavior. For example, advances in optics and intracellular calcium (Ca^2+^) indicators have allowed for simultaneous stimulation and observation of individual neurons within animals ^1–3^. In parallel, a wave of new genetically encoded tools has further advanced our ability to monitor and perturb neural activity ^4–8^. Though these technological advancements are promising, they are limited in accessibility and application to the field of neuroscience as a whole. The cost, both in terms of finance and human resources, of new optical microscopic systems is a barrier that many labs are unable to overcome. A solution to this problem may lie in recently developed genetically engineered tools which do not require large optical configurations to modulate and interact with.

Genetic technologies have often relied on pharmacological agents for temporal specificity. Due to slow pharmacological dynamics and long half-life, the temporal resolution at which pharmacologically based tools can window activity-dependent gene transcription severely limits their use in behavioral research. Recently, tools have been developed to enable researchers to gate gene expression in neurons by dynamic changes in the concentrations of calcium (Ca^2+^) within the cell bodies of individual neurons, known as cytosolic Ca^2+ 9,10^. While localized transduction of cytosolic Ca^2+^ can be used as a biomarker of long-term cellular changes in individual neurons, the inherent ambiguity of divergent cytosolic Ca^2+^-signaling pathways and their relationship to extracellular signaling greatly limits our ability to interpret the meaning behind them ^11–15^. For example, it is unclear if a given Ca^2+^ signal is due to a subthreshold oscillation, a train of action potentials, or release of an intracellular Ca^2+^ store ^12,16–20^. Additionally, these technologies are composed in large genetic encodings which greatly hinders their practical use in *in vivo* behavioral experiments. To be effective, such technologies will need to possess key characteristics including: (i) behaviorally relevant temporal dynamics of its gating mechanisms, (ii) minimal cross-reaction with other cellular molecules, and (iii) ease of use in a variety of organisms. We are particularly interested in detecting calcium signals that regulate activity-dependent gene expression important for learning and memory, as well as for the propagation of hyperactivity of neurons that occurs during epileptogenesis and neurodegeneration.

Several studies have shown nuclear Ca^2+^ transients evoked by experience-driven synaptic activity to be distinct events representing the propagation of information mediated by action potentials ^21–26^. We set out to make a tool that would detect such Ca^2+^ transients using a genetic readout relevant to endogenous activity-dependent gene regulation. This tool, called CLiCK (Ca^2+^ Light Coincidence Knock_in/out_), is a dual condition genetic expression system. In neurons, CLiCK acts as a coincidence detector for the co-occurrence of neural activity (as monitored by the presence of transient nuclear Ca^2+^) and blue light (∼450 nm). Transcription occurs only through the simultaneous release of both gating mechanisms. Neither light nor activity alone are sufficient to induce reporter gene expression. We demonstrate high temporal responsiveness of CLiCK *in vivo*, enabling subsecond windowing of behaviorally-driven neural activation, in contrast to pharmacological windows that last many hours. To eliminate failures due to incomplete delivery of transgenes using multiple viral vectors, we engineered the entirety of CLiCK into a single multicistronic unit < 2.5 KB; small enough to deliver via a single AAV vector and containing a transcriptional marker (ZsGreen) for vector detection.

The photo-switch in CLiCK comes from the light-oxygen-voltage (LOV) sensitive domain from *Avena sativa* (AsLOV_2_) ^27,28^. The C-terminus of native AsLOV_2_ contains a photo-reactive Jα-helix domain. In the absence of light, the Jα-helix adducts into core β-sheets of the phototropin, burying several hydrophobic residues. Exposure to blue light induces core structural changes, originating at the flavin mononucleotide (FMN) binding pocket, which propagate to the C-terminus causing the Jα-helix to unfold, exposing its residues. The photo-driven abduction and thermal reversal of LOV domains establish photo-stationary equilibrium in the sub millisecond time scale ^29,30^, well positioned to capture a behaviorally relevant event. In addition to thermal reversion, UV light can be used to drive LOV domains back to their adducted ground state at rates of picoseconds ^31^.

To make the CLiCK transcription factor (CLiCK_tf_), we linked an NLS-Gal_4_ DNA binding domain to the N-terminus of the phototropin AsLOV_2_ using a flexible GS-linker (to decrease the chances of functional interference between protein domains). Independent of light, CLiCK_tf_ localizes to its specific DNA binding site, the upstream activator sequence (UAS), positioned upstream of the mRuby_2_ transgene (**Fig. 1a**). A chimeric Jα-transactivator domain (AD) (see below for identification) was engineered into the C-terminus for light-switchable allosteric gating to attract co-activator proteins (**Fig. 1b**). In the dark state, catalytic residues of the AD are buried in the core “cage” of the protein. When exposed to blue light, conformational changes within the core sheets of the transcription factor propagate to the C-terminus. This propagation leads to unfolding of the Jα-helix, subsequently exposing critical residues of AD that are recognized by transcriptional cofactors necessary to drive transcription of mRuby_2_.

**Figure 1.**
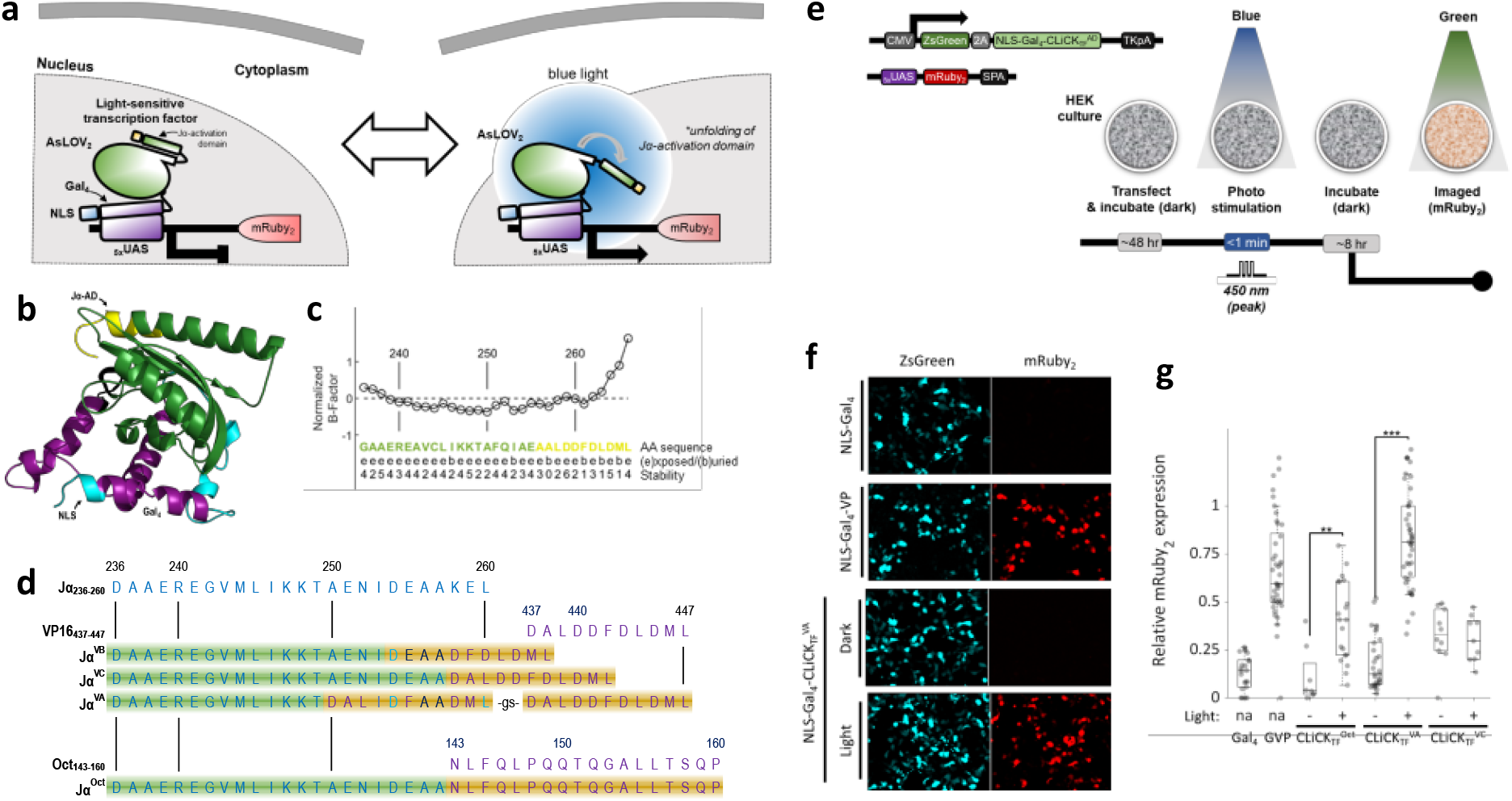
Engineering a light-responsive transcription factor: CLiCK. (**a**) Schematic of light-gated transcription factor, CLiCK_tf_. In dark conditions, CLiCK_tf_ acts to gate gene expression through sterically blocking key catalytic residues along its transcriptional activation domain (Jα-AD). Exposure to blue light induces core conformational changes within Jα-AD which lead to the unfolding and exposure of its transcriptional activation domain, enabling interactions with transcriptional co-factors, as indicated by the arrow on lower right. NLS, nuclear localization sequence; AD, activation domain. (**b**) Example ribbon model of CLiCK_tf-iVP16E_ in low-energy dark state as determined by iterative template fragment assembly simulations. An NLS-Gal_4_-domain (cyan-purple) is linked by a flexible GS linker to the N-terminus of the phototropin AsLOV_2_ (green) from *Avena sativa*. At its C-terminus, a chimeric Jα-transactivation domain is contained (yellow). Photo-sensitive chimers were evaluated *in silico* from their amino acid sequences. Predicted characteristics of depict favorable dark-state packing (solvent accessibility = 2) and stability (normalized B-Factor = -0.15) of presumptive critical residue F261. Normalized β-Factor values below 0 indicate stable residuals; predicted solvent accessibility range from 0 (highly (b)uried, shown by letter “b” in figure) to 9 (highly (e)xposed, shown by letter “e” in figure). (**d**) The Jα-domain amino acid sequence of wild-type AsLOV_2 (236-260)_ and the minimal activation motifs of VP16 _(435-447)_ and Oct _(143-160)_ showing the four Jα-AD permutations originally selected from *in silico* models for testing *in vitro*. Color of residue letters: Jα, blue; transactivators VP16 or Oct, purple; chimeric, cyan. Small-case depicts linker. Background coloring indicates Jα region and putative transactivation domain. (**e**) Schematic representation of the fluorescence reporter system for testing CLiCK_tf_ light-gating *in vitro*. (top) The diagram illustrates the constitutively expressed fluorescence transfection marker (ZsGreen) expression cassette that contains the transgene for a CLiCK_tf_, NLS-Gal_4_ (negative control) or NLS-Gal_4_-VP16 (positive control) and its corresponding _5x_UAS target in the fluorescence functional reporter (mRuby_2_) vector. (bottom). 2A; self-cleavage peptide. Dissociated HEK293 cells were co-transfected with a CLiCK_tf_ plasmid and its corresponding functional reporter plasmid. Following a period of incubation, cells were imaged for mRuby_2_ (green excitation light) then either briefly exposed to blue light or returned to light-tight containers without exposure to blue light. After 6 hours cells were imaged for mRuby_2_. When not being transfected, imaged, or photo-activated, cells were kept in light-tight containers. (**f**) Example images of CLiCK_tf_^VA^ light-responsive expression of mRuby_2_. Top row shows negative control of mRuby_2_ expression under transcription factor with activation domain removed (NLS-Gal_4_). Top middle row shows positive control of mRuby_2_ expression under a light-independent transcription factor (NLS-Gal_4_-VP). Bottom middle row shows of mRuby_2_ expression under photo-sensitive transcription factor (NLS-Gal_4_-CLiCK_tf_^VA^) in absence of light. Bottom row shows of mRuby_2_ expression under photo-sensitive transcription factor (NLS-Gal_4_-CLiCK_tf_^VA^) following brief photo-stimulation. (**g**) Relative mRuby_2_ expression from imaging data plotted with boxplot superimposed. The three different CLiCK_tf_ chimeras showed differential light-responsive expression of mRuby_2_ where CLiCK_tf_^VA^ had the greatest dynamic range. CLiCK^tf Oct^: 0.11 ± 0.15, n = 861 (-Light), 0.40 ± 0.21, n = 2184 (+Light), P = 0.001 - 0.01; CLiCK ^tf VA^: 0.19 ± 0.14, n = 1811 (-Light), 0.83 ± 0.25, n = 2397 (+Light), P < 0.001. Boxes show median line, 25^th^ and 75^th^ percentiles edges and min/max outlier whiskers. Two sample T-tests were performed. ***p<0.001, ** p = 0.001 - 0.01, *p = 0.01 - 0.05. Reported values are the means, standard deviations, and cell counts for each condition. CMV: cytomegalovirus, 2A: 2A self-cleaving peptide, NLS: nuclear localization sequence, Gal_4_: Gal_4_ DNA binding domain, TKpA: tymidine kinase polyadenylation, UAS: upstream activation sequence, SPA: synthetic polyadenylation.

To identify ADs suitable for caging within AsLOV_2_, we searched for candidates that were biochemically similar to the Jα-helix and two transactivator peptides were chosen: the virion protein 16 of herpes simplex virus type 1 (VP16) and the lymphocyte-derived octamer transcription factor 2A (Oct). The amphipathic α-helix of VP16 contains a minimal activation motif (VP16_437-447_) DALDDFDLDML capable of inducing expression in either proximal or remote locations ^32^. VP16_437-447_ negative residues D443 and D445 are critical for initial docking interactions while hydrophobic residues L439, F442, and L444 are critical for stable transcriptional activity with residue F442 particularly critical ^33^. The Oct minimal activation motif (Oct_143-160_) NLFQLPQQTQGALLTSQP induces proximal transcriptional activation ^32,34^. As mutations deep inside the Jα-helix are more likely to disrupt adduction into the β -sheets of AsLOV_2_, we designed preliminary Jα-AD permutations which focused towards the outer segment of the Jα-helix. Those permutations conserved hydrophobic residues I539, A542, and A543 known to make critical contacts with AsLOV_2_ domain β-sheets ^35,36^.

The I-TASSER protein modeling suite ^37^ was used to construct high quality models to identify favorable Jα-AD chimeric sequences (**Fig. 1b**). For each chimeric sequence an average of 9,869 simulations was performed, generating a large ensemble of structural conformations from which five full length atomic models were constructed. Structural and functional analyses were performed using the models with the greatest confidence score. Impacts of AsLOV_2_-Jα alterations on stability of Jα-AD segments, particularly along key hydrophobic residues, were evaluated for impact on gating integrity (**Fig. 1c**). General stability of Jα-ADs was determined by averaging predicted inherent thermal mobility (B-factor) of its residues. Additionally, solvent accessibility of individual residues along Jα-ADs was examined to determine if particular residues packed favorably against the protein core. Finally, to assess for any fundamental disruption of AsLOV_2_ photo-transduction capability, the ability of chimeric sequences to bind FMN was examined using protein-protein interaction methods COFACTOR ^38^ and COACH. Chimeric models were benchmarked against a model of wildtype AsLOV_2_^404-546^ (^wt^AsLOV_2_).

Robust confidence scores of chimera models ranged from 0.83 to 0.97 (μ = 0.88). Estimated root-mean-square deviation (RMSD) ranged from 2.9Å to 3.1Å (μ = 3.0Å), falling within previously established resolutions for functionally predictive models. All models were found to maintain high confidence (μ = 0.97) of FMN binding at their corresponding C450 binding pocket of ^wt^AsLOV_2_, suggesting compatibility of Jα-AD sequences to dock with core β-sheets. Predicted β-factors (BFP) for Jα-ADs ranged from -0.3 to 0.1, where BFP values higher than 0 are less stable, and were found comparable to ^wt^AsLOV_2_ (BFP_Jα_ = 0.14). Chimeric sequences showing favorable dark-state packing and stability of critical residues were genetically engineered for *in vitro* assays (**Fig. 1d**).

To evaluate the gating properties of each chimera, a photo-activation assay was performed in human embryonic kidney (HEK293) cells (**Fig. 1e**). Cells were co-transfected with two plasmids: 1) an NLS-Gal_4_-CLiCK_tf_ chimera placed under a constitutive promoter, and 2) a _5x_UAS-mRuby_2_ target reporter with a transfection marker (ZsGreen). For positive controls, an NLS-Gal_4_-VP16_tf_ replaced NLS-Gal_4_-CLiCK_tf_. For negative controls, VP16_tf_ was removed from the positive construct. Following transfection, cells were kept in light-tight containers. To induce light-dependent gene expression, cells were briefly exposed to 1 second pulses of blue light (450 nm peak, 12 mW). Following photo-stimulation, cells were immediately returned to light-tight containers where they remained until imaging. All experiments were carried out in light-controlled environments.

With the exception of CLiCK_tf_^VB^ (data not shown), all CLiCK_tf_ chimeras were capable of driving mRuby_2_ expression. Only CLiCK_tf_^Oct^ and CLiCK_tf_^VA^ showed light-dependent dynamic expression (**Fig. 1g**). CLiCK_tf_^VA^ produced a particularly strong response to light, a 6-fold increase in expression compared to NLS-Gal_4_-Jα^VA^. Additionally, CLiCK_tf_^VA^ dark-state expression did not differ from NLS-Gal_4_ negative controls, ∼12% to 14% respectively.

We exploited the endogenous signaling pathway of intranuclear DREAM (downstream responsive element antagonist modulator) ^39,40^ as a sensor for _in_Ca^2+^ transients that are associated with activity-dependent gene expression (**Fig. 2a**). In neurons, DREAM functions as a principal transcriptional switch regulating the on/off status of specific activity-dependent gene expression programs that control synaptic plasticity, and learning and memory ^41–44^. At basal [_in_Ca^2+^], DREAM binds to a downstream regulator element (DRE) within promoter regions of the genome. Upon rising [_in_Ca^2+^], Ca2^+^ binds directly to DREAM inducing its dissociation from the DRE ^39,45–48^. The conserved, orientation-independent ^41^, sequence “GTCA” forms the core of DRE motifs while flanking nucleotides determine DREAM binding affinity ^49,50^. TATTTTGG<u>ACTG</u>G GTA (iDRE), from the inducible cAMP early repressor gene (ICER), has previously been shown to bind tightly to DREAM ^49^. To minimize Ca^2+^-independent expression of the reporter, iDRE was incorporated into the DNA vector between the UAS and mRuby_2_ so that expression could be gated by both light and nuclear Ca^2+^ (**Fig. 2b**). In addition to iDRE, we tested GGAGTCAGC (cDRE) from the presumptively high-affinity cross-species prodynorphin consensus DREs, PuNGTCAPuPuG ^50^.

**Figure 2.**
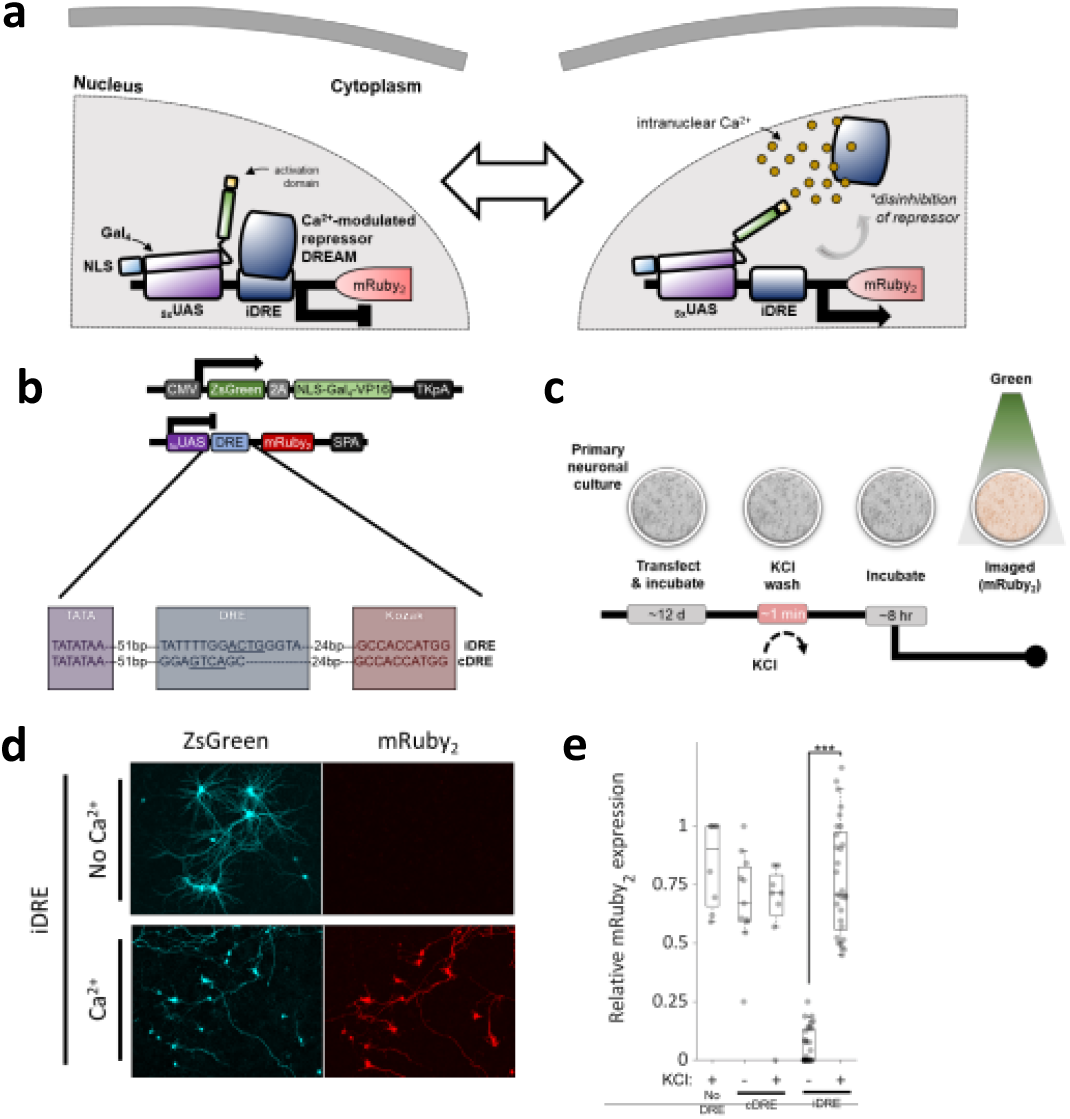
Engineering of CLiCK’s intranuclear Ca^2+^-gate. (**a**) Schematic overview of intranuclear Ca^2+^-gated expression system. At basal intranuclear Ca^2+^ (_in_Ca^2+^), DREAM represses downstream gene expression. Ion fluxes of _in_Ca^2+^ cause dissociation of DREAM from its binding sequence through direct Ca^2+^ interactions, relieving transcriptional repression. (**b**) Schematic representation of constitutively expressed transactivator (top) and its downstream _in_Ca^2+^-dependent fluorescence reporter (middle) containing either an iDRE or cDRE motif (bottom). (**c**) *In vitro* neuronal assay design for activity-induced _in_Ca^2+^-dependent expression of mRuby_2_. Dissociated E15 rat primary cortical neurons in culture were co-transfected with a vector set depicted in and incubated for ∼12 days. To induced activity driven intranuclear Ca^2+^ transients, 20 mM of KCl was applied to the culture media and rapidly washed out with preconditioned media (see Methods). Immediately following a KCl wash, cells were returned to the incubator for 6-8 hours before imaging for expression of mRuby_2_. (**d**) Example images of CLiCK_act_^iDRE^ showing _in_Ca^2+^-gated expression of mRuby_2_ for KCl induce neural activity (bottom) and TTX treat control (top). (**e**) Relative mRuby_2_ expression from imaging data plotted with boxplot superimposed. No DRE: (+) 0.84 ± 0.18, n = 137; cDRE: (-) 0.69 ± 0.2, n = 236 and (+) 0.64 ± 0.27, n = 43; iDRE: (-) 0.06 ± 0.08, n = 654 and (+) 0.78 ± 0.23, n = 1818. Boxes show median line, 25^th^ and 75^th^ percentiles edges and min/max outlier whiskers. Two sample T-tests were performed. ***p<0.001, ** p = 0.001 - 0.01, *p = 0.01 - 0.05. Reported values are the means, standard deviations, and cell counts for each condition.

All vector sets containing the dual DNA sensors express mRuby_2_ in the absence of DREAM, as assessed by transfection assays in HEK cells which do not express endogenous DREAM protein. To evaluate Ca^2+^-dependent gating capacities of iDRE and cDRE within the new mRuby_2_ vector, we used a conditional activity-induced *in vitro* neural assay (**Fig. 2c**). Primary neuronal cultures were co-transfected with CMV-NLS-Gal_4_-VP16-2A-ZsGreen (where 2A is a self cleaving peptide sequence) and a vector containing a _5x_UAS-DRE-mRuby_2_ reporter with either the iDRE or cDRE sequence positioned between the UAS and mRuby_2_. For a positive control, neurons were transfected with the Ca^2+^-independent _5x_UAS-mRuby_2_ reporter. While cDRE was ineffective at repressing mRuby_2_ expression in absence of _in_Ca^2+^, iDRE tightly regulated expression with basal levels of 6% compared to 78% in the absence of DREAM, a 13-fold dynamic range (**Fig. 2e**). Moving forward, the iDRE cassette was engineered into the dual Ca^2+^-light gated expression system called CLiCK.

CLiCK contains a photo-sensitive transactivator (CLiCK_tf_) and a direct _in_Ca^2+^-regulated expression cassette (CLiCK_act_). These two gates operate in tandem, simultaneously gating gene expression (**Fig. 3a**). KCl induced-activity and light exposure was combined to determine whether coincidence gating of the fluorescent mRuby_2_ reporter could be detected in primary neuronal cultures under the following four conditions: inactive-dark, inactive-light, active-dark, and active-light (**Fig. 3b**). Briefly, to evaluate baseline expression levels of mRuby_2_ in the co-absence of both light and activity (inactive-dark), cells were kept in light-tight containers and treated with 1 μM of TTX to suppress spontaneous neural activity. Baseline leak of expression through each gate was examined by individually applying either the KCl or light assay as described in Figures 1e and 2c with the supplemental treatment of TTX to block spontaneous neural activity in the light assay (**see Methods**). Reporter expression (mRuby_2_) under the co-occurrence of activity and light (active-light) was examined by the simultaneous application of KCl and light protocols. To temporally define the window of activity upon exposure to light, culture media was immediately exchanged with TTX (1 μM) conditioned media following all KCl washes. Immediately following all conditions, cells were returned to light-tight containers and incubated for 6 hrs before imaging. With relief of both Ca^2+^ and light gates, mRuby_2_ expression increased by 19-fold compared to expression in the absence of activity and light (**Fig. 3d**). When both gates are engaged, we observed minimal expression (transcriptional leakage) of mRuby_2_ (mean of 3.9%). With the release of either gate, leak increased slightly where 5.8% is observed through the Ca^2+^ gate and 12.9% through the light gate. Though transcriptional leakage is more prominent in active cells under dark conditions, it is comparable to baseline observed in HEK cells (**Fig. 1g**).

**Figure 3.**
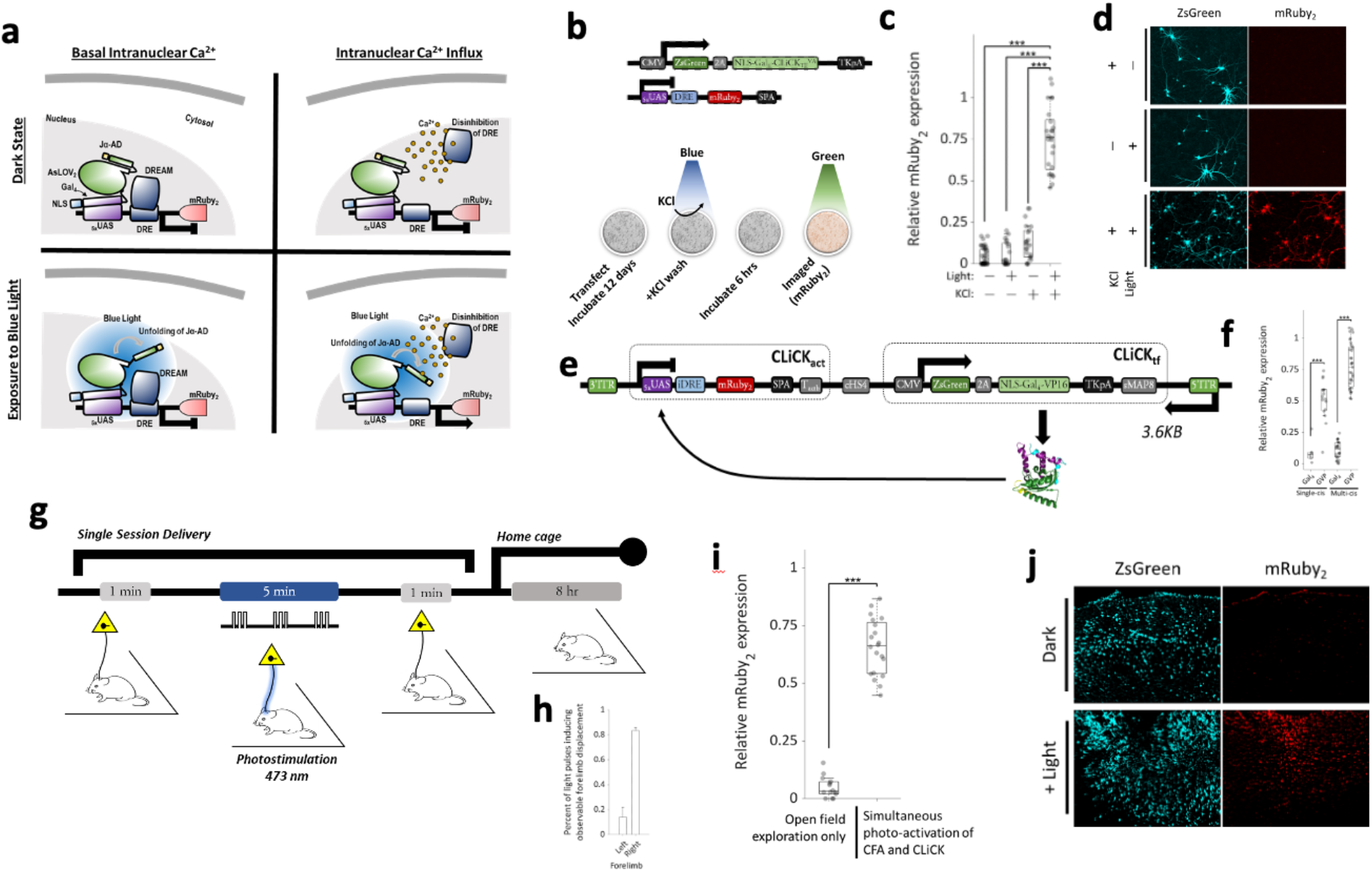
Evidence for _in_Ca^2+^-light coincident expression and a platform for *in vivo* optimization and future use. (**a**) Schematic of CLiCK. Combining gates depicted in Figures 1a and 2a, CLiCK gates gene expression through two independent gating-mechanisms: (1) an intranuclear Ca^2+^-gated repressor and (2) a light-gated transcription factor. In the absence of either blue light or activity-induced _in_Ca^2+^, transcription of the mRuby_2_ transgene is blocked. The mRuby_2_ reporter is only detected upon the co-occurrence of activity-induced increases in _in_Ca^2+^ and the presence of blue light. Neither alone is enough to induce its gene expression. (**b**) Schematic representation of the (top) CLiCK fluorescence reporter system and (bottom) *in vitro* neuronal assay design used for light and activity coincidence expression of mRuby_2_. (**c**) Relative mRuby_2_ expression from imaging data plotted with boxplot superimposed. (-/-) no stimuli: 0.04 ± 0.05, n = 1221, P < 0.001; (+/-) induced-activity only: 0.13 ± 0.10, n = 990, P < 0.001; (-/+) blue light only: 0.06 ± 0.07, n = 591, P < 0.001; (+/+) induced-activity and blue light: 0.75 ± 0.19, n = 838, P < 0.001. (**d**) Example images of CLiCK coincidence gating of mRuby_2_. (**e**) Schematic circuit design for single vector CLiCK. Consists of two expression cassettes, CLiCK_act_ and CLiCK_tf_. A gene-of-interest can be easily inserted in-frame via AgeI and NotI restriction sites within CLiCK_ACT_. The insulator cHS4 is inserted between expression cassettes, and a transcriptional pause sequence (T_act_ and sMAP8) was inserted near the end of each expression cassette. Length of entire vector is 3626 bp (without reporter) and 4340 bp with reporter mRuby_2_ as conditional reporter transgene. (**f**) Relative mRuby_2_ expression differences between multi-plasmid and single-plasmid vector designs of CLiCK. To characterize the dynamic ranges of mRuby_2_ expression between single-vector and multi-vector designs, a NLS-Gal_4_-VP (labelled as GVP in the figure) transcription factor was used to evaluate peak expression. To characterize minimal expression the activation domain VP_16_ was removed from the transcription factor (labelled as Gal_4_ in the figure). Relative expression was as follows: Single-cis: 0.09 ± 0.09, n = 442 (Gal_4_), 0.51 ± 0.16, n = 1335 (GVP), P < 0.001; multi-cis: 0.11 ± 0.06, n = 2218 (Gal_4_), 0.78 ± 0.17, n = 3915 (GVP), P < 0.001. (**g**) Schematic for testing CLiCK *in vivo*. An AAV_2/9_ virus encoding the entirety of CLiCK (including the transduction marker ZsGreen and conditional reporter mRuby_2_) was injected bilaterally into the caudal forelimb area (CFA) of the motor cortex of Thy1-ChR2 transgenic mice. To induce activity in only the right forelimb, a single optical fiber was implanted in the left hemisphere targeting the CFA. For photo-induction, a 473 nm light was delivered to the left CFA at 9.6 mW (measured at pre-implanted fiber tip). Light pulses of 10 Hz (100 ms period) were delivered at 50% duty cycle (50 ms ON, 50 ms OFF) for one second. In total, 50 light pulses (5 sec interpulse intervals) were delivered per session. Sessions lasted a total of 5 mins. Following stimulation, mice were returned to their home cage for 8 hrs then perfused for imaging analysis. (**h**) Summary of behavioral data. Average number of light-induced forelimb displacements observed per session (left 5 mice; right 4 mice). (**i**) Summary of cell count results in postmortem tissue after behavioral experiment. In absence of light (right hemisphere), exploration of an open field did not induce expression of mRuby_2_ in the CFA. Delivering light pulses (left hemisphere) during exploration of an open field induced behavioral activation of the contralateral forelimb and expression of mRuby_2_. Open field only: 0.03 ± 0.04 (n = 1120 cells/5 mice); co-activation: 0.66 ± 0.12 (n = 1479 cells/4 mice), P < 0.001. (**j**) Representative brain sections from experiment in (g) for non-light activated (top) and light co-activated ChR and CLiCK (bottom). Left hemisphere received photostimulation protocol, whereas right hemisphere was kept in the dark. Simultaneous light induced CFA activity and CLiCK drives expression of mRuby_2_ in the light condition (bottom) but not in the dark condition (top). Boxes show median line, 25th and 75th percentiles edges and min/max outlier whiskers. Two sample T-tests were performed. ***p<0.001, ** p = 0.001 - 0.01, *p = 0.01 - 0.05. Reported values are the means, standard deviations, and cell counts for each condition.

To prepare CLiCK for practical use in experimental settings, the entirety of CLiCK (< 2.5 KB) was engineered into a single multicistronic AAV vector (**Fig. 3e**) leaving, minimally, 2.3 KB for insertion of a conditional transgene. In its concise form, CLiCK contains two expression cassettes: (1) CLiCK_act_ and (2) CLiCK_tf_. The promoter region of CLiCK_act_ contains an iDRE binding sequence 51 bp downstream of a _5x_UAS sequence and 24 bp upstream from the Kozak sequence recognized by the ribosome. Conditional transgenes can be easily inserted in-frame via AgeI and NotI restriction sites. CLiCK_tf_ contains genes that encode a transduction marker (ZsGreen) and the photo-switchable transcription factor CLiCK_tf_, separated by a P2A self cleaving peptide sequence. The CLiCK_tf_ gene was placed downstream of the P2A to avoid addition of undesired residues left behind by cleavage. To enhance stable transcription termination and avoid conditional expression leaks arising from runoff transcription, CLiCK_act_ was placed upstream of CLiCK_tf_ and a pause sequence (Tact and sMAP8, respectively) was inserted near the end of each expression cassette ^51^. Expression variability through unintended enhancer activity or position effects of transgenes, was minimized using an insulator sequence (cHS4) inserted between expression cassettes. Changes in mRuby_2_ expression under positive conditions were evaluated in HEK cells using a Gal_4_-VP16 transcription factor (**Fig. 1g**). For negative controls, the VP16 activation domain was removed such that only the Gal_4_ DNA binding domain remained (**see Methods for details**). Compared to delivery of CLiCK via multiple vectors, we found positive expression of mRuby_2_ under a single multicistronic vector increased by 28%, indicating improved delivery of the complete package of transgenes (**Fig. 3f**).

Most importantly, the potential *in vivo* application of CLiCK was tested using Thy1-ChR2-YFP mice (n = 5) (Jackson Labs; B6.Cg-Tg(Thy1-ChR2/EYFP)18Gfng/J), a mouse line that has been engineered to express channel rhodopsin 2 under control of the thymus antigen 1 (Thy1) promoter. The caudal forelimb area (CFA) of the mouse motor cortex was bilaterally injected with a recombinant AAV containing CLiCK (**Fig. 3g**). A fiber optic targeting the CFA was unilaterally implanted in the left hemisphere of each mouse such that photo-activation of the left CFA would produce displacement of the right forelimb. The right hemisphere was treated as a dark control. Following 3 weeks post-surgery, mice were placed in an open field (91 × 91 cm) and a brief photo-stimulation protocol was implemented (**for details see Methods**). For photo-induction, a 473 nm light was used which activates both ChR2 and CLiCK’s photo-switchable transcription factor. A light pulse of 10 Hz (100 ms period) was delivered at 50% duty cycle (50 ms ON, 50 ms OFF) for one second. In total, 50 light pulses (5 sec interpulse interval) were delivered per mouse. Sessions lasted a total of 5 mins. Following photo-stimulation, mice were returned to their home cage for 8 hrs then perfused for imaging analysis. On average, photo-stimulation of the left CFA induced an observable right forelimb displacement for 83% of the light pulses delivered in a session (**Fig. 3h**). Without photo-stimulation, free mobility of the forelimbs in open field exploration produced < 5% expression leak of mRuby_2_ in CFA cells (n = 1120 cells/5 mice) (**Fig. 3i**). With the co-occurrence of light and forelimb activity, there was a 13-fold increase of mRuby_2_ expression in the CFA (n = 1479 cells/4 mice).

We have identified and addressed three crucial characteristics currently limiting the potential of current technologies. These include a (i) poor temporal resolution in the temporal windowing of signal (activity) -dependent gene transcription, (ii) signal ambiguity through divergent signal transduction pathways, and (iii) impractically complex packaging of genetic material. Collectively, these limitations present a very real barrier between the application and potential of optogenetic tools.

To address limitations in the temporal resolution of physiologically conditioned (activity-dependent) gene transcription we engineered a light-sensitive transcription factor which minimizes both the number of molecular events and physical distance between the site of light transduction and the site of gene transcription to within the nucleus of the cell. To dispel ambiguity in the signal transduction of neural activity we incorporated a direct nuclear Ca^2+^- modulated repressor upstream of our reporter gene. Finally, through concise genetic design and encoding we are able to package and deliver the entirety of our system within a single AAV vector, thus overcoming practical barriers in its application *in vivo*. Together, these features offer a potential platform technology for the practical application in studies of neural activity patterns.

In our *in vivo* proof of concept study, we used light to drive both neural activity (via channelrhodopsin) and the unfolding of CLiCK’s light-sensitive transcription factor. This allowed us to link a behavioral readout (displaced limb movements) to anatomically specific expression of mRuby_2_ in the CFA of M1. With the combined vector, we observed that transcription of the reporter mRuby_2_ corresponded with a decreased expression of the transduction marker, ZsGreen (**Fig. 3j**). This may be due to a competition in the molecular co-factors required for transcription of nearby genes, especially with increased activation of one promoter, or epigenetic modification of the CMV promoter driving the transduction marker. When we delivered CLiCK via two vectors (3b top) we observed continuous expression of both the ZsGreen and mRuby_2_ with the co-occurrence of light and KCl induced Ca^2+^ (3d).

As this technology continues to be develop, I am particularly interested in exploring its applications in translational and clinical neuroscience to address unmet medical needs in mental health conditions. I believe there are several points along the development of a therapeutic candidate which precise experimental control of activity-dependent gene transcription would be useful, if not groundbreaking. These include, but not limited to, the development of new neurocircuit-based animal disease models, behaviorally relevant (functional) transcriptomics, and behaviorally-targeted delivery of new gene therapy candidates.

## Methods

### In Silico Modeling

To construct high-quality predictions of 3D protein structure and function from amino acid sequences an *I*terative *T*hreading *ASSE*mbly *R*efinement (I-TASSER) algorithm was adopted. The I-TASSER algorithm consists of 3 consecutive steps of threading, fragment assembly, and iteration ^52–54^. 3D models were visualized and plotted using the Python package for molecular visualization, PyMOL.

### Molecular Cloning

#### Photo-switchable chimerics

To create the functional domains (DNA binding domain (Gal_4_), photo-sensitive core (AsLOV_2_), and transcription activation domain (AD) of CLiCK_tf_, we incorporated and synthesized DNA fragments from multiple sources. For the light sensitive core of the CLiCK_tf_, the LOV protein was used in combination with other elements. The expression construct for *Avena sativa* phot1 LOV2 (Uniport O49003_AVESA) was generously provided by Dr. Andreas Möglich (Humboldt University of Berlin) in a ET-28c plasmid. The Gal_4_ DNA binding domain and UAS promoter sequence was generously provided by Dr. Ben Wolozin (Boston University) in a pCI-neo plasmid. Single-stranded NLS and Jα-activation domain oligonucleotides were synthesized using Integrated DNA Technologies custom DNA service. Oligos ≤ 60 nucleotides were purified using standard desalting purification while oligos > 60 nucleotides were purified using PAGE. Oligos were resuspended in nucleus-free buffer and annealed at equal molar concentrations. For in-frame cloning of Jα-activation domains into the N-terminus of AsLOV_2_, overlap extension PCR ^55^ was used. AsLOV-Jα-AD PCR products were then cloned in-frame into pGal_4_-Cl backbones using standard digest and ligation protocols. For Gal_4_-Jα-AD constructs, double stranded Jα-AD oligos were used to clone into the pGal_4_-CI.

#### Direct intranuclear Ca^2+^-dependent repressor of expression

Another component of the system, CLiCK_act_, was engineered to repress gene transcription in a intranuclear Ca^2+^-dependent manner. To engineer the functional domains of CLiCK_act_ (UAS promoter region, DNA downstream regulatory element (DRE), reporter gene (mRuby2)) we incorporated and synthesized DNA fragments from multiple sources. pmRuby_2_-N1 plasmid was obtained from Addgene (accession # U55762). Single-stranded TATA box and DRE oligonucleotides were synthesized using Integrated DNA Technologies custom DNA service. Oligos were resuspended in nucleus-free buffer and annealed at equal molar concentrations. Standard PCR, digest and ligation protocols were used for subcloning double-stranded TATA, UAS and DRE fragments into pmRuby_2_-N1.

#### Rational optimization for in vivo applications

The expression construct for LVDP ^51^ was generously provided by Dr. Stelios T. Andreadis (University at Buffalo, Amherst, NY) in a pCS vector (Addgene plasmid # 12158).

### Cell Cultures

#### HEK cultures

To provide an initial *in vitro* test system, we used cultured HEK cells. HEK cells were plated in 24-well, 35mm or 100mm dishes such that their confluency was approximately 60% on the day of transfection. Cells were transfected with Lipofectamine 2000 (Thermofisher).

#### Primary neuronal cultures

To provide an in vitro test system with neurons, we created primary neuronal cultures. Primary hippocampal and neocortical neurons were dissected from embryonic day 18 (E18) Sprague-Dawley rats (Charles River Laboratories). Pregnant dams were euthanized by exposure to 95% CO2 and embryos were harvested. Embryonic brains were removed and placed in ice-cold Ca^2+^/Mg^2+^ free (CMF) media [Ca^2+^/Mg^2+^ free Hank’s BSS, 4.2 mM sodium bicarbonate, 1 mM pyruvate, 20 mM HEPES, 3 mg/ml BSA, pH7.25-7.3]. Relevant brain regions were dissected, homogenized, and centrifuged in plating media [Neurobasal media (Invitrogen), 10% fetal bovine serum (Gibco), 100 U/ml penicillin, 100 μg/ml streptomycin, 200 mM glutamine]. Neurons were plated on poly-L-lysine (0.1 mg/mL) coated dishes, placed in an incubator (37°C/ 5% CO2) for one h, after which plating medium was removed and replaced with defined medium [Neurobasal media (Invitrogen), B27 serum-free supplement (Gibco), 100 U/ml penicillin, 100 μg/ml streptomycin, 200 mM glutamine]. Cultures were returned to the incubator for approximately 10 days until further use for testing expression and combined activation of reporter by light and calcium activation.

### Transfections

#### HEK transfections

The following procedures were used in order to transfect the HEK cells with our reporter systems. Plasmid DNA and Lipofectamine (1:3 ratio) were individually diluted in reduced-serum Opti-MEM media, mixed gently and incubated for 5 min at room temperature. Following incubation, dilutes were combined, mixed gently and incubated for 20 minutes at room temperature. After 20 mins of incubation, DNA-Lipofectamine complexes were added to HEK media in a 1:5 complex-to-media ratio, briefly mixed on a rocker plate and then incubated at 37°C in a 5% CO2 incubator for 24-48 hours prior to testing for transgene expression. For light-sensitive experiments, cells were kept in light-tight containers following transfection.

#### Primary neuronal transfections

The following procedures were used to introduce the genetic components of our systemin cells of neuronal cultures. NeuroMag transfection was performed using primary neuronal cultures at 7 DIV, using the NeuroMag Magnetofection™ kit (Oz Biosciences #KC30800). Plasmid DNA was diluted in NBM and added to the NeuroMag Transfection Reagent. The DNA solution was incubated (20 minutes, room temperature) before being added to neuronal culture dishes and incubated on a magnetic plate (20 min, room temperature) provided with the kit. Cells were removed from the plate and incubated (37°C, 5% CO2) for 10-14 days before assaying.

### Cell Assays

#### Photo-switchable cell assay

To induce light-dependent transcription, cells were briefly exposed to 1 second pulses of blue light (450 nm peak, 12 mW). Following photo-stimulation, cells were immediately returned to light-tight containers and placed into a cell incubator for 6-8 hours prior to imaging. All light sensitive experiments were carried out in a light-controlled environment.

*KCl induced neuronal activity assay*. For activity-dependent transcription conditions, intranuclear Ca^2+^ transients were induced by applying 20 mM of KCl to neuronal culture media and rapidly washed out using neuronal culture media drawn from the sample 3-7 days prior to assaying. Following KCl washes, cells were returned to the incubator for 6-8 hours before imaging. To suppress transcription through spontaneous neural activity in negative control conditions, 1 μM of TTX was add to the culture media 6-8 hours prior to imaging.

#### Photo-switchable activity-dependent assay

KCl induced-activity and light assay designs were combined and coincidence gating of the fluorescent reporter mRuby_2_ in primary neuron cultures was examined under four conditions: inactive-dark, inactive-light, active-dark, and active-light. To evaluate leakage of mRuby_2_ expression in the co-absence of both iCa^2+^and light (inactive-dark), cells were kept in light-tight containers and treated with 1 μM of TTX to suppress spontaneous neural activity. Independent leak of expression through each gate was examined by individually applying either the iCa^2+^ or light assay (previously described above) with the supplemental treatment of 1μM TTX to culture media to block spontaneous neural activity in the light-alone assay. For co-occurrence of iCa^2+^ and light (active-light), simultaneous application of KCl and light protocols was performed. Culture media was immediately exchanged with 1 μM TTX preconditioned media following all KCl washes to create a temporally windowed period of KCl-induced neural activity. Immediately following all light-sensitive conditions, cells were returned to light-tight containers and incubated for 6-8 hrs before imaging.

### In Vitro Image Acquisition, Processing, and Analysis

#### Image Acquisition

Fluorescent images were collected on an Olympus inverted fluorescence microscope at 10X and 20X using a mercury bulb for excitation of fluorophores. ZsGreen was excited at 470/40 nm (center wavelength/bandwidth) and emissions were collected at 525/50 nm, emission (495 nm dichroic mirror). eYFP was excited at 500/20 nm and emissions were collected at 535/30 nm, emission (515 nm dichroic mirror). mRuby2 was excited at 560/40 nm and emissions were collected at 630/60 nm, emission (585 nm dichroic mirror).

#### Image Processing and Analysis

Automated cell counts were performed in ImageJ using a custom written script. Briefly, images were first convolved with a Gaussian filter with a standard deviation of 2. To avoid erroneous segmentation of non-fluorescent images, background subtractions were performed on images with an intensity distribution kurtosis > 0.4 using the Subtract background plugin with a rollsize 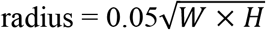 pixels, where W = image width and H = image height. Additionally, to automate appropriate thresholding for segmentation, images with an intensity distribution kurtosis > 0.4 were checked for in mean/median ratio ≥ 1 and skewness > 0.1. For images meeting all three criteria, the Triangle ^56^ threshold was used, otherwise, the Moments ^57^ threshold was used. Next, binary masks were created from thresholds and an erosion and dilation was performed on masks followed by a Watershed segmentation. Particles with an area greater than 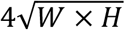 pixels squared were the counted using the Analyze Particles plugin.

### Mice

For rapid photo-induced motor behavior, experiments were performed using a transgenic mouse line that expresses the light-activated ion channel, Channelrhodopsin-2, fused to Yellow Fluorescent Protein, in layer 5 cortical neurons under control of the mouse thymus cell antigen 1 (Thy1) promoter (B6.Cg-Tg(Thy1-ChR2/EYFP)18Gfng/J, Jackson Labs). Male and female mice (2-4 months old) were housed by gender in Plexiglas cages together with their siblings before surgery, but separated for individual housing after surgery, and maintained on a 12 h light/dark cycle. Food and water was provided *ad libitum* throughout the duration of housing.

### Stereotactic Surgeries

#### General surgical procedure

All procedures were done in accordance with the Boston University Institutional Animal Care and Use Committee. Surgeries were performed to inject virus containing the CLiCK construct. Viral packaging of CLiCK was performed at Boston Children’s Hospital Viral Core. In preparation for surgery, animals were given an injection of atropine (10 mg/kg IP) and buprenorphine (0.1 mg/kg IP) and anesthetized with isoflurane (3% induction, 1-2% during surgery). Animals were given buprenorphine (0.1 mg/kg SC), ketofen (2-5 mg/kg SC) and baytril (10 mg/kg SC) during a 5d postsurgical care period and given 3 weeks for viral transduction and full recovery before beginning of photo-activation.

#### Viral transduction

During surgery, the caudal forelimb areas (CFA) (1.0AP, 1.5ML, 0.6DV) of the motor cortex in mice were bilaterally injected with an AAV containing CLiCK. A craniotomy was performed 1 mm anterior and 1.5 mm lateral to bregma, and the injection needle (34 g, beveled; WPI) was lowered 0.6 mm at a 0° polar angle, following stereotactic coordinates from Paxinos and Franklin (2008). A total of 250 nL of the pAAV_2/9_-Gal_4_-mRuby-cHS4-CMV-ZsGreen:2A:CLiCK_TF_ (titer = 1.95E+13 GC/mL; 0.05 M PBS) was infused at a rate of 100 nl/min (UMP3 electrical pump, WPI). Once the full viral injection was delivered the injection needle was left in place for 10 minutes to prevent backflow of the injected virus solution.

#### Fiber optic implants

Following viral injections, a mono fiber-optic cannula (200 μm core, NA: 0.48; MFC_200/230-0.48_2.5mm_SMR_FLT, Doric Lenses) targeting the CFA (1.0 AP, 1.5 ML, 0.5 DV) was chronically implanted in the left hemisphere such that photo-activation of the left CFA would produce displacement of the right forelimb. The right hemisphere was treated as a dark control. Three anchoring screws were positioned across the skull and, along with the fiber-optic implant, were cemented onto the animal’s skull.

### In Vivo Experimental Outline

#### Open-field environment

Photo-stimulation was carried out as mice explored an open-field environment (1 × 1 m) with 60 cm high black walls. The behavior of the mouse before, during and after laser stimulation of channelrhodopsin was recorded overhead at 30 frames per second (fps) using a 12.2 megapixel CMOS video camera (Sony Exmor IMX363). Photo-induced limb displacement during laser stimulation was defined by the co-occurrence of the photo-stimulation and rapid pulsing displacement of the right forelimb due to activation of the left motor cortex.

#### Photo-stimulation

For simultaneous laser stimulation of ChR2 and CLiCK_tf_ in mice, a fiber-coupled 473 nm laser (OBIS FP, Coherent), was digitally modulated via the TTL channel with an Arduino Due microcontroller using a custom built script (see Appendix 2). Laser light was delivered into the brain via a system of fiber patch cords (Thorlabs) and a rotary joint (FRJ_11_FC-FC, Doric Lenses). The last connection was a magnetic connection to the implanted light fiber. The laser light power entering the implanted light fiber was measured before and after every recording session and adjusted before the recording session to yield an estimate of 9.6 mW laser power delivered into the CFA. Laser stimulation was delivered at 10 Hz (100 ms period) with a 50% duty cycle (50 ms ON, 50 ms OFF) for one second with 5 s interpulse intervals (a typical session lasted 300 s with 51 Laser OFF and 50 Laser ON periods).

### Histology

Following photo-stimulation, mice were returned to their home cage for 8 h. Mice were then deeply anesthetized by isoflurane or intraperitoneal injection of Euthasol (390 mg/kg), and subsequent transcardial perfusion with saline followed by 10% buffered formalin (SF100-4, ThermoFisher Scientific). Brains were extracted and post-fixed in formalin for 2-4 more days after which they were placed in a 30% sucrose solution in KPBS for 1-2 additional days. The brains were then frozen and sliced on a cryostat (Leica CM 3050S) in 40 μm sections after which they were mounted and coverslipped with Vectashield Hardset mounting medium (Vector Laboratories). Slices were then imaged at 4x, 10x, and 20x on a Nikon Eclipse Ni-E epifluorescence microscope to verify proper placement of the fiber optic fibers above the CFA of the motor cortex.

### In Vivo Image Acquistion, Processing, and Analysis

#### Image Acquisition in Post-Mortem Tissue After In Vivo Photo-Activation

The effect of in vivo activation on the expression of fluorescence protein was analyzed with post-mortem imaging of the histological preparations of the tissue. Fluorescent images of the prepared tissue were collected on a Nikon Exclipse Ni-E epifluorescence microscope at 10X (Plan Fluor 10x NA 0.3) and 20X (Plan Apo Lambda 20x NA 0.75) using a Sola Light Engine for excitation of fluorophores. ZsGreen was excited at 470/40 nm (peak/bandwidth) and emissions were collected at 525/50 nm, emission (495 nm dichroic mirror). eYFP was excited at 500/20 nm and emissions were collected at 535/30 nm, emission (515 nm dichroic mirror). mRuby_2_ was excited at 560/40 nm and emissions were collected at 630/60 nm, emission (585 nm dichroic mirror).

#### Image Processing and Analysis of post-mortem histological preparations

Automated cell counts of fluorescence labelled cells from histological preparation were performed in ImageJ using a custom written script (see Appendix 1) as described in the Methods section

## Acknowledgements

We would like to acknowledge the mentorship and support of the Michael Hasselmo laboratory during the course of this technology development and the members of the Russek and Hasselmo laboratories, with especial thanks to Sabita Bandyopadhyay, Zhuting Li, and Kathryn Hixson.

## Notes

This work was supported by NINDS 5R01NS051710-14 and the BU Provost award for collaborative neuroscience research.

### Competing Interest Statement

The authors have declared no competing interest.

